# Visual reconstruction from 2-photon calcium imaging suggests linear readout properties of neurons in mouse primary visual cortex

**DOI:** 10.1101/300392

**Authors:** Stef Garasto, Anil A. Bharath, Simon R. Schultz

**Affiliations:** Department of Bioengineering, Imperial College London; Centre for Neurotechnology, Imperial College London

## Abstract

Deciphering the neural code, that is interpreting the responses of sensory neurons from the perspective of a downstream population, is an important step towards understanding how the brain processes sensory stimulation. While previous work has focused on classification algorithms to identify the most likely stimulus label in a predefined set of categories, fewer studies have approached a full stimulus reconstruction task. Outstanding questions revolve around the type of algorithm that is most suited to decoding (i.e. full reconstruction, in the context of this study), especially in the presence of strong encoding non-linearities, and the possible role of pairwise correlations. We present, here, the first pixel-by-pixel reconstruction of a complex natural stimulus from 2-photon calcium imaging responses of mouse primary visual cortex (V1). We decoded the activity of approximately 100 neurons from layer 2/3 using an optimal linear estimator and an artificial neural network. We also investigated how much accuracy is lost in this decoding operation when ignoring pairwise neural correlations. We found that a simple linear estimator is sufficient to extract relevant stimulus features from the neural responses, and that it was not significantly outperformed by a non-linear decoding algorithm. The importance of pairwise correlations for reconstruction accuracy was also limited. The results of this study suggest that, conditional on the spatial and temporal limits of the recording technique, V1 neurons display linear readout properties, with low information content in the joint distribution of their activity.

## Introduction

Neural firing patterns in primary sensory areas are commonly thought to contain information about the external world. For example, neurons in the primary visual cortex (V1) respond preferentially to specific properties of visual stimuli, such as orientation and location [1]. To understand how such information is encoded in the neural responses and how it is used to drive behaviour, it is crucial to investigate how it can be extracted (decoded) from the the firing patterns of populations of neurons [2, 3]. Indeed, this would give us insights into how a downstream population could perform the same task [4] and which aspects of the neural code are essential for good decoding performance [5–7]. Furthermore, neural decoding algorithms have important practical implications for brain-machine interface devices [8, 9].

Decoding techniques have been applied to many species and brain areas, mostly to classify neural response patterns according to a predefined set of labels [10–12].

However, fine stimulus details also contribute to shape our visual perception, and it appears plausible for this to be reflected in the response pattern of many brain regions, especially at the early stages of the visual stream [13]. How much information about these fine details can be recovered from neural activity in V1? Given that the neural encoding process is highly non-linear, do we need strong non-linearities in the decoding algorithm (hereafter intended as full stimulus reconstruction) as well to achieve optimal performance? So far, previous work has provided inconclusive evidence. Botella-Soler et al. [14] showed that a non-linear decoder outperformed a linear one when reconstructing complex movies from retinal ganglion cells, and similar results have been reported by others [15–17]. In contrast, some studies achieved near optimal performance using a linear decoder [2, 18–20]. Here, we address these questions through pixel-wise reconstruction of natural scenes from 2-photon calcium imaging in mouse V1. We intend to investigate what are the requirements for an algorithm capable of extracting such information.

To the best of our knowledge, full visual stimulus reconstruction has never been attempted with responses from mouse V1, potentially because of the low acuity of mouse vision. However, given the importance that the mice have gained in recent years as a model for mammalian vision [21, 22], building a baseline of performance and establishing the feasibility of full visual stimulus reconstruction could help further the understanding of the neural coding of vision in mice, as well as other animals. At the same time, much effort has been given to reconstructing stimuli using a wide variety of recording techniques and experimental conditions. Bialek et al. first estimated a one dimensional stimulus from single neuron recording in the fly visual system [2]. Subsequently, other noteworthy examples, in addition to the ones previously mentioned, include the reconstruction of natural movies from cat LGN using liner spatio-temporal filters [23], and geometrical shape estimation from monkey V1 using deconvolution with a point spread function [13]. Thirion et al. extended the field to artificial visual scene reconstruction from fMRI recording in humans [24]. Naselaris et al. [25] and Nishimoto et al. [26] later reconstructed natural scenes and movies, respectively, and showed that performance improved when using a large, but fixed, dataset of natural images as the prior. More recently, Wen et al. [27] used a deep neural network as the common underlying model for both the encoding and decoding inference, while Parthasarathy et al. [17] decoded simulated retinal ganglion cells responses by enhancing linear reconstructions with a specific type of neural network architecture, namely a convolutional autoencoder.

A common thread among most of the cited studies is population decoding, which is beneficial because it allows not only the reconstruction of more complex stimuli, but also the investigation of how the interaction between multiple neurons influences the decoding process [28]. The ability of two-photon imaging to scale up to hundreds and even thousands of cells [29, 30] makes it an appealing recording technique when the aim is to decode from large groups of neurons recorded simultaneously. It is, thus, essential to investigate benefits and limitations of using 2-photon imaging for stimulus reconstruction. Finally, we would argue that natural stimulation is better suited to studies on stimulus reconstruction that aim at getting insights into how downstream populations might solve a similar task. Indeed, since neural response properties tend to vary under different types of stimuli [31], using artificial images might, in principle, lead to different results [23].

Here, we set out to reconstruct visual scenes from 2-photon imaging recordings of a neural population in mouse V1 [32]. We first use the optimal linear estimator (OLE) [19], and then investigate the impact of pairwise correlations by building a correlation-blind linear decoder (dOLE) [6, 28, 33]. Then, we compare the accuracy of the linear reconstruction with that obtained using a feedforward artificial neural network (ANN) [34]. We report that a simple linear decoder can rival an artificial neural network in extracting information about pixel intensities from neural activity in mouse V1.

## Results

We reconstructed visual stimuli from 2-photon imaging recordings of mouse V1. The data was released under a Creative Common Attribution License by Antolik et al. [32]. The input of the reconstruction algorithm is the total spike count of a population of 103 neurons during the presentation of each frame, while the output is the pixel-wise reconstruction of the visual stimulus. Neural responses were standardised to 0 mean and unit standard deviation, while the pixel values were all between 0 and 1.

### A simple linear decoder can reconstruct natural images from mouse V1

First, we used an optimal linear estimator (OLE) to reconstruct the visual stimuli. Despite its simplicity, a linear decoder has been shown to be able to achieve good performance [18, 19, 23] and is compatible with highly non-linear encoding mechanisms [4]. The optimal linear decoding filters were computed to minimize the mean squared error (MSE) of the reconstruction, subject to L_2_ regularization. The relative weight of the regularization term was computed using cross-validation. Performance was quantified using the MSE and the Pearson correlation coefficient (*ρ*) between each target and reconstructed frame and taking their median values over all frames in the test dataset. The full distributions are shown in Fig 1A and 1B: the median value was 〈*MSE_f_*〉 = 0.069 for the MSE, and 〈*ρ_f_*〉 = 0.283 for the correlation coefficient. Some of the frames are predicted with good accuracy, although the distributions are spread across a broad range of values, with some reconstructed frames also being anti-correlated with the targets.

**Fig 1.**
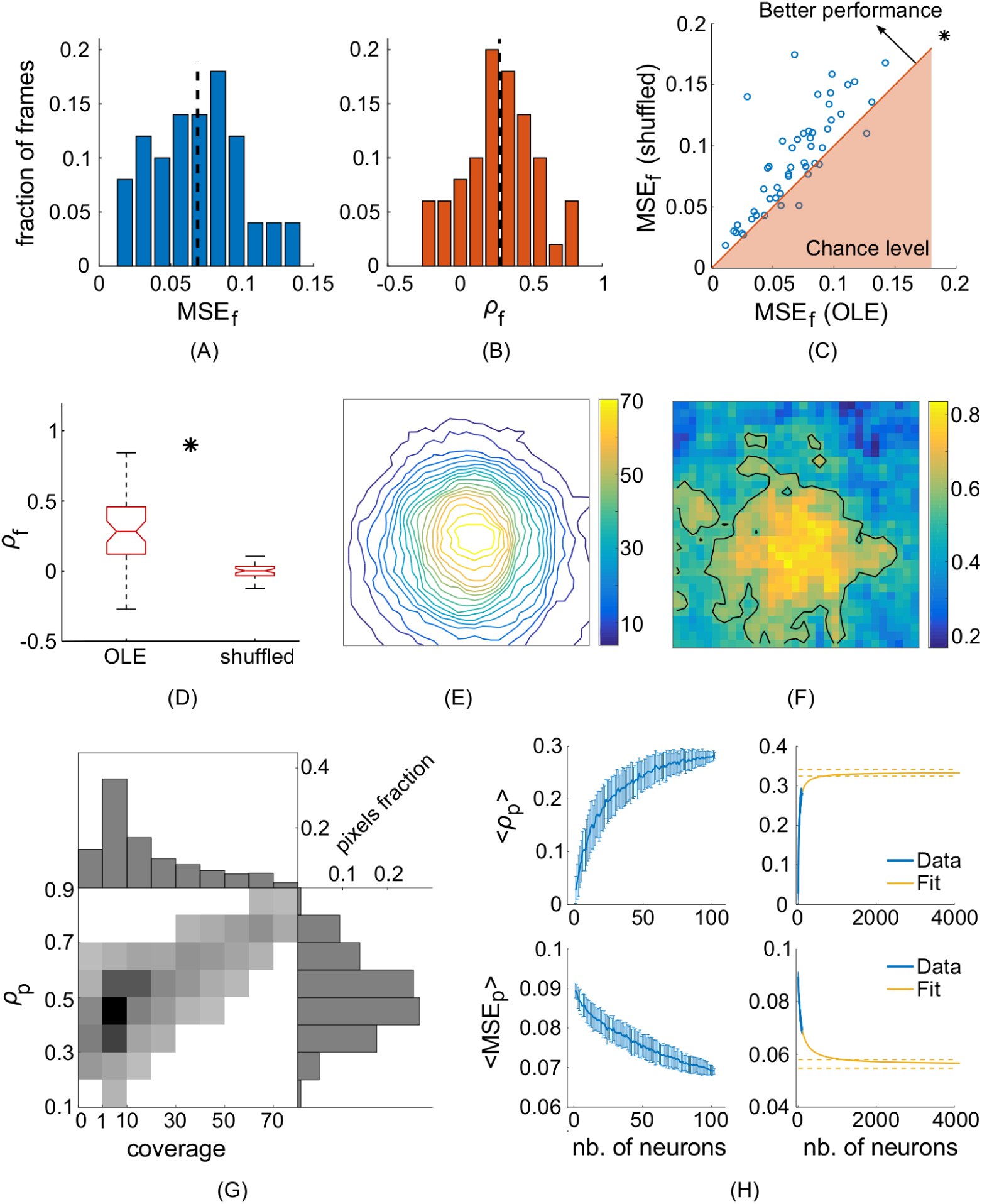
Linear decoder performance. **(A)** Distribution of the MSE values between each target and linearly reconstructed frame in the test dataset. The black dotted line represents the median value. **(B)** Same as (A), but with the distribution of the frame-by-frame correlation coefficient *ρ_f_*. **(C)** Performance (MSE) of the non-linear decoder across frames, compared with chance level performance. The asterisk implies that the OLE is significantly better than chance (p<0.001). **(D)** Same as (C) but with the frame-by-frame correlation coefficient. **(E)** Receptive field coverage level across the visual field of view. The contour lines indicates how many receptive fields include each individual pixel. **(F)** Performance of the linear decoder across pixels, in terms of correlation coefficient *ρ_p_* computed between the target and reconstructed pixel intensity profiles. **(G)** Pixel-wise performance of the linear decoder vs receptive field coverage. **(H)** Median correlation across frames as a function of the number of neurons used in the decoder. Blue dots and bars are, respectively, the means and standard deviations across various equally sized subsets of neurons. The yellow solid and dotted lines are, respectively, the extrapolated performance curve and the confidence bounds for the asymptote value.

To evaluate whether the OLE performed better than chance, we used a neural shuffling procedure to destroy the input-output relationship between the responses and the stimuli. Briefly, each spatial pattern of responses in the training dataset was randomly assigned to a different stimulus frame, while the identity of each neuron was preserved (non preserving the identity gave the same results), then the OLE was trained on these shuffled data and tested on non shuffled responses in the test dataset. This procedure was repeated 200 times: the mean MSE and correlation coefficient for each frame over the repeated shuffles gave a distribution that represents the chance level of performance for the OLE (the control distributions). To test the null hypothesis of chance level performance, A Wilcoxon-signed rank test (AB test 1) was performed between the control distributions and those obtained from non-shuffled neural responses (Fig 1B). The *p*-values for both the MSE and the correlation coefficient were lower than 10^−8^, meaning that the OLE performance are significantly better than chance. For completeness, a second statistical test was performed to assess the likelihood of obtaining the median performance values on the test dataset purely given by chance (AB test 2, more details can be found in the methods): *p*-values for both the MSE and the correlation coefficient were lower than 10^−16^.

We found a strong positive relationship between the performance of the algorithm at each individual pixel, and the number of receptive fields that included that pixel (Fig 1D,1E). The level of coverage was computed by fitting a linear receptive field on the training data using the linearised version of the receptive fields obtained after fitting each neuron with a pyramid wavelet model (Fig 1C). This is likely due to the increase in information content about an individual pixel that derives from having more neurons being stimulated by that pixel. Fig 1D also shows that the better predicted pixels cluster in the middle part of the image, consistently with the fact that the stimulus was centered around the location of the population receptive fields. These considerations imply that increasing the number of neurons recorded might further increase the performance of the algorithm by allowing for better coverage of a greater number of pixels, especially those that are further away from the center. However, Fig 1f seems to indicate otherwise, since it only predicts an increase of 17.9%, 18.6%, for the correlation coefficient and the MSE, respectively. The asymptote values and confidence intervals for the two performance measures are 0.33 ± 0.01 and 0.0564 ± 0.0016, respectively. This can be explained by the fact that the extrapolation procedure we used might not be able to accurately estimate the increase in performance we would obtain by imaging neurons whose receptive fields are shifted towards the sides of the field of view. Instead, it likely makes predictions based on a proportional increase in coverage. This is less than optimal, since the central pixels are already close to saturation. Nonetheless, the performance improvement predicted with this method could be considered a lower bound on the expected gain derived from decoding a larger population of neurons. Finally, the reconstructed frames (49 out of 50, in total) are shown in Fig 2. Overall, the linear decoder seems to perform better on large bright and dark patches than on the smaller, high resolution details.

**Fig 2.**
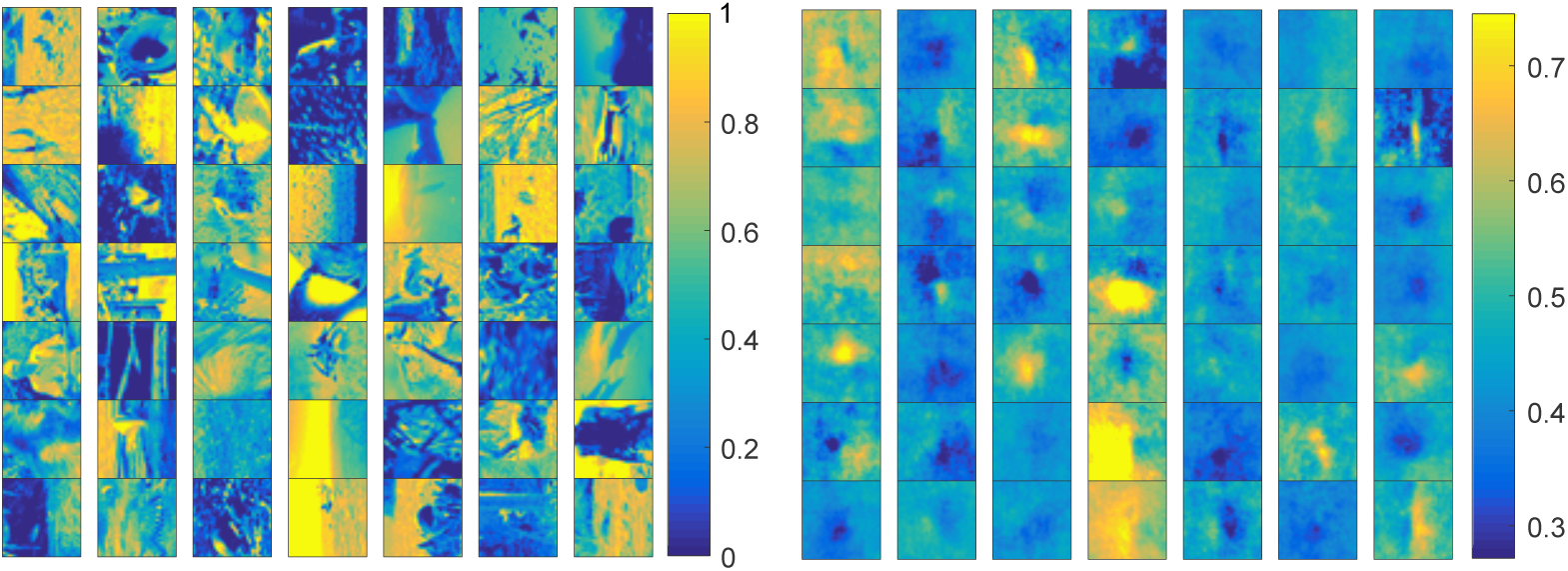
Linear reconstruction results. Target images are shown on the left, and the reconstructed ones on the right, presented in the same order (49 frames out of 50 total are shown). Each group of images has its own colourmap.

Finally, it is worth noting that one of the limitations of the linear decoder is that each pixel is reconstructed independently from the others. That is, it is not specifically designed for multi-target regression and correlated output. To check that this did not impact reconstruction performance, we tested another linear algorithms, specifically the Multi-target Regressor Stacking (details can be found in the S1 Appendix caption). However, this algorithm did not perform better than the naive OLE.

### Removing pairwise correlations has a limited effect on linear decoding performance

Neurons often show at least some degree of correlated activity, meaning that their spiking behaviour is not independent from one another [6]. However, the role of correlated activity in the context of stimulus decoding is still not fully understood [6, 28]. The implications are of both theoretical and practical importance. Indeed, if neurons can be considered as independent encoders, then the problem of decoding from a population is equivalent to that of combining the outcome of multiple individual decoders, which is technically and computationally easier [35]. To investigate this aspect of the neural code, we built a correlation-blind linear decoder [6, 28, 33] by setting to zero all the off-diagonal elements of the matrix **R^*T*^R**, which carry the full extent of the information about pairwise correlations between neurons. We named this decoder the diagonal Optimal Linear Estimator, or dOLE. It is worth noting, though, that this approach removes both signal correlations, the covariation in the mean response of pairs of neurons across stimuli, and noise correlations, the trial-to-trial variability [36]. Indeed, given that the training dataset is single-trial, both types of correlations are present, but it is not possible to separate them.

The results of applying the dOLE in comparison with the performance of the OLE are shown in Fig 3, where each point represents how well an individual frame has been reconstructed using the two algorithms (OLE on the y-axis and dOLE on the x-axis). It can be seen that the data points cluster around the *y* = *x* line, meaning that there is not a substantial difference in performance between the OLE and the dOLE. There are, however, few frames that seem particularly affected by the removal of correlations. To quantify whether there is any significant difference between the performance of the two algorithms, we compared the two distributions using a Wilcoxon-signed rank test against the null hypothesis of OLE being at most as good as the dOLE. The resulting *p*-values for the MSE and the correlation coefficient are 0.13 and 0.048, respectively, showing that the difference in performance are not that statistically significant. This could suggest that, in this case, neurons might be considered as independent encoders. However, it could also depend on other factors, such as a relatively low amount of pairwise correlations in this specific dataset (so that the OLE itself depended only marginally on the off-diagonal elements), or the small size of the test dataset.

**Fig 3.**
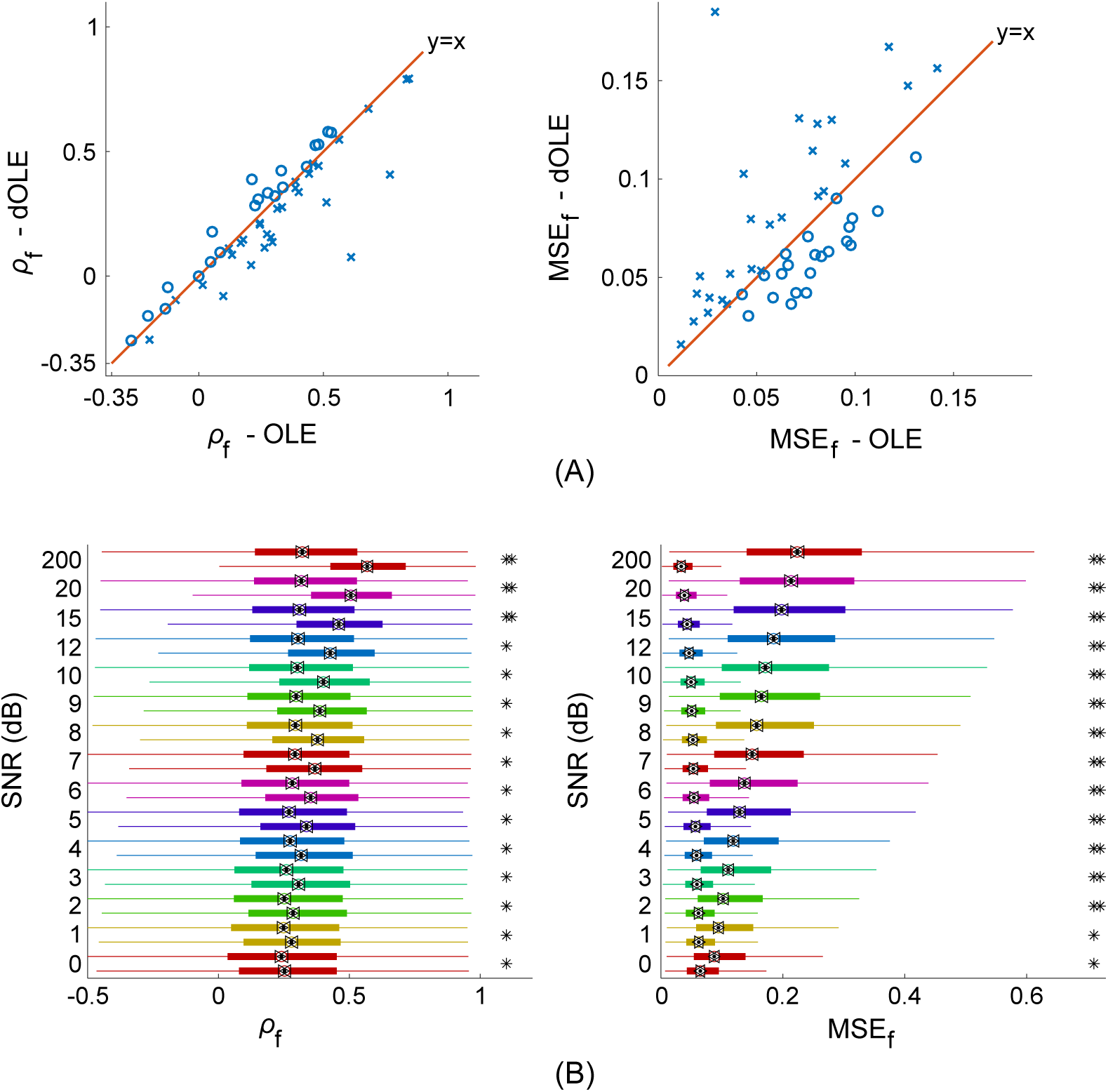
Linear decoding with and without pairwise neural correlations. **(A)** Comparison of distributions of performance across frames between OLE (y axis) and dOLE (x axis), when decoding from recorded data. **(B)** Comparison of distributions of performance across frames between OLE (y axis) and dOLE (x axis), when decoding from surrogate data at various SNR levels. The presence of an asterisk implies that the OLE is significantly better than the dOLE (p<0.001). The second asterisk means that the same is true when comparing groups of 50 frames.

Hence, we performed some further analysis on synthetic data generated after fitting a pyramidal wavelet model to the recorded training dataset [25, 26, 32, 37]. Briefly, we created a bank of Gabor wavelets at different scales, locations, frequencies and phases and transformed the stimulus from pixel space to wavelet space using a cascade of linear and non-linear operations (half-wave rectification and sum of squares of quadrature phases wavelets were used as the non-linearities of the model). The median correlation coefficient of the model fit across neurons was 0.43. Then, we generated synthetic data using a separate, and much larger (approximately 18000 samples for training, and 3500 for testing), dataset of natural images, that has been contrast-normalised to match the distribution of the original dataset. For the noise model, we added uncorrelated Gaussian noise at different levels of SNR. By construction, this surrogate dataset only contains signal and not noise correlations.

The results of applying both the OLE and the dOLE at different SNRs are shown in Fig 3B, where each pair of same colour box plots represent the performance distributions of the two algorithms (the OLE below and the dOLE above) for the same noise level. The same Wilcoxon-signed rank test was used to assess statistical significance (with Holm-Bonferroni correction to control for multiple comparisons): the presence of at least one asterisk on the right means that the obtained p-value was less than 0.01. To better compare results with recorded data, we repeated the same Wilcoxon-signed rank test on 100 random subset of 50 frames: the presence of a second asterisk on the right of each box plot pair means that such a test was significant at *α* = 0.01 at least 99 times out of 100. Therefore, the presence of two asterisks implies that the differences between the OLE and the dOLE are robust with respect to the number of frames in the test dataset.

Overall, it can be observed that, on surrogate data, the OLE performs much better than the dOLE, independently on the noise level, which would suggest that weak pairwise correlations do not necessarily result in a lack of differences between the two algorithms. Furthermore, in some cases, and indeed most of the times with respect to the MSE, this results do not depend on the amount of samples in the test dataset, implying that differences between OLE and dOLE could be evident also on the small test dataset available for the experimental data.

### Non-linear reconstruction using a neural network

Despite the performance of the OLE being significantly better than chance, it is worth asking whether a more powerful decoder would be able to achieve better reconstruction results. Such a question is relevant to not only quantify how much information a downstream population would be able to extract from the neural responses, but also which algorithmic strategy would be more successful. Hence, we investigated whether relaxing the linearity assumption of the OLE was able to increase the accuracy of the visual stimulus reconstruction. To this end, we built a feed-forward Artificial Neural Network (ANN) that takes as input the neural responses and return the predicted frames as the output. Such a network was trained end-to-end using backpropagation. While theoretical results proved that a multi-layer ANN is capable of approximating any function, in practice it can often be challenging to train the network properly: a sufficient amount of data and a careful selection of the architecture, the loss function and the hyper-parameters are needed. Here, we analysed a large number of different combinations and selected the best by comparing performance either on the training data or on a separate validation dataset (approximately 10% of the training samples). Some results for the hyper-parameters space exploration can be found in the supporting material (S2 Fig). The final model is a two-layers fully connected ANN, with 64 and 32 units in the first and second layer, respectively; *tanh* as the unit non-linearity; batch normalisation and dropout (with probability 0.5 and 0.4 for first and second hidden layer, respectively) to regularise the training and prevent overfitting. Furthermore, the neural responses were normalised to 0 mean and unit standard deviation, and binary cross-entropy was used as the loss function. Finally, we trained the network using ADAdelta, and found that 200 epochs were usually enough to reach convergence, with the learning rate starting at 1 (the default value for ADAdelta) and then decreased to 0.001 after 100 epochs. To improve our chances of selecting a better local minimum, we trained 54 different initialisations of the model and chose the one with the highest performance (in terms of median correlation coefficient) on a validation dataset. Ensemble performance is shown in panel A of S3 Fig. Results for the reconstructions obtained with the ANN are shown in Fig 4A. The median values for the correlation coefficient and the MSE are 0.33 and 0.054, respectively. Both seem marginally better than those obtained with the OLE. To assess the significance of the results, we repeated the neural shuffling tests already described. The *p*-values obtained were lower than 10^−4^ for AB test 1 and lower than 10^−16^ for AB test 2.

**Fig 4.**
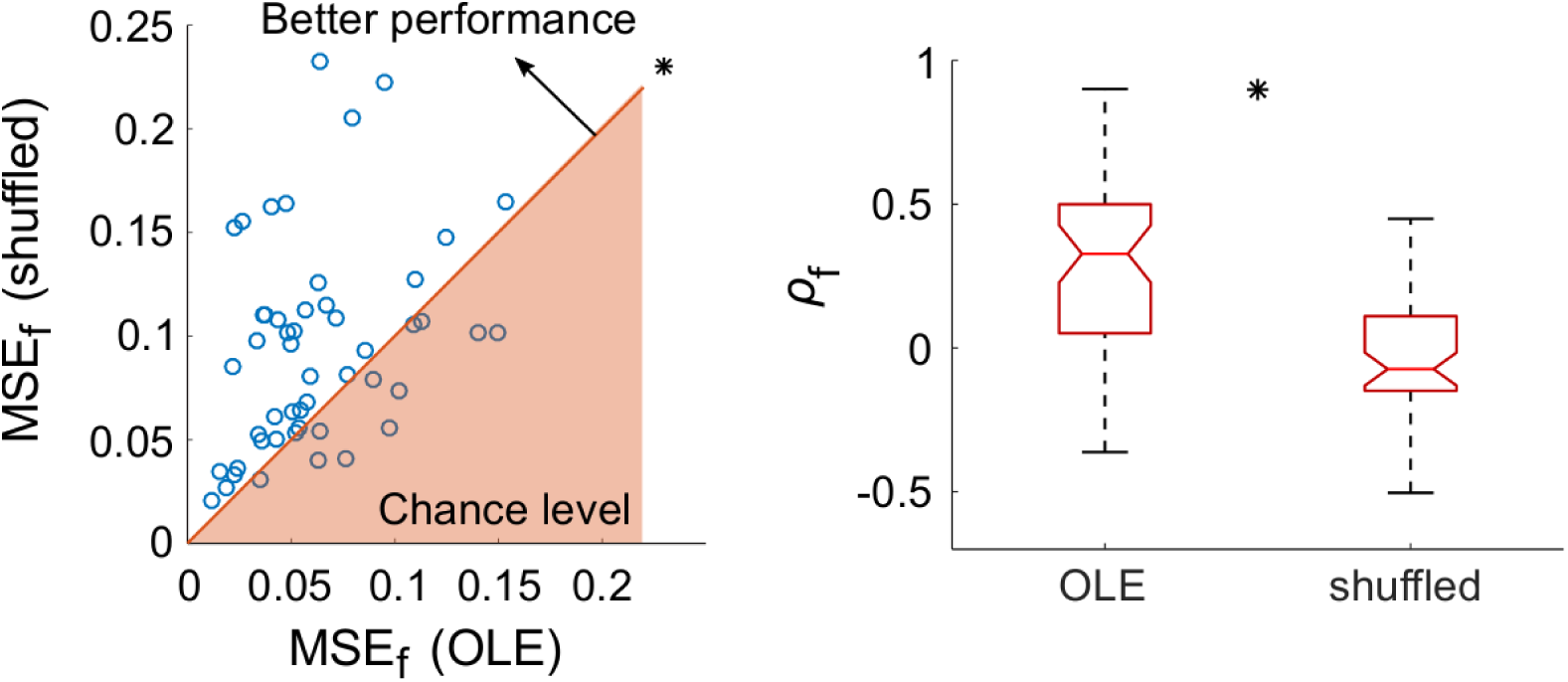
Non-linear decoding: accuracy. Performance of the non-linear decoder across frames, for both MSE (left) and correlation coefficient (right), compared with chance level performance. The asterisk implies that the OLE is significantly better than chance (p<0.001).

Some example reconstructions are presented in panel B of S3 Fig, while Fig 5 shows some statistics representing what the network has learnt and how it operates. Specifically, we visualised the weights connecting the second layer units to the output units (Fig 5C). Such weights can be interpreted as 32 different 31 × 31 matrices, one for each node in the second hidden layer and the reconstructed stimuli can be seen as produced by a weighted combination of such matrices. The influence of each basis on the final output is proportional to the activation of the respective hidden unit. Hence, visualising the weights of the output layer can give some insight into the type of reconstructed stimuli we can expect from the network. In this case, it appears that there is a prevalence of low frequency features that are likely to characterise the final reconstructions as well, even though some higher resolution details might still arise from their weighted combination. It is also worth noting that the weights appear to be spatially smooth, even though no explicit L_2_ regularisation was used in the output layer.

**Fig 5.**
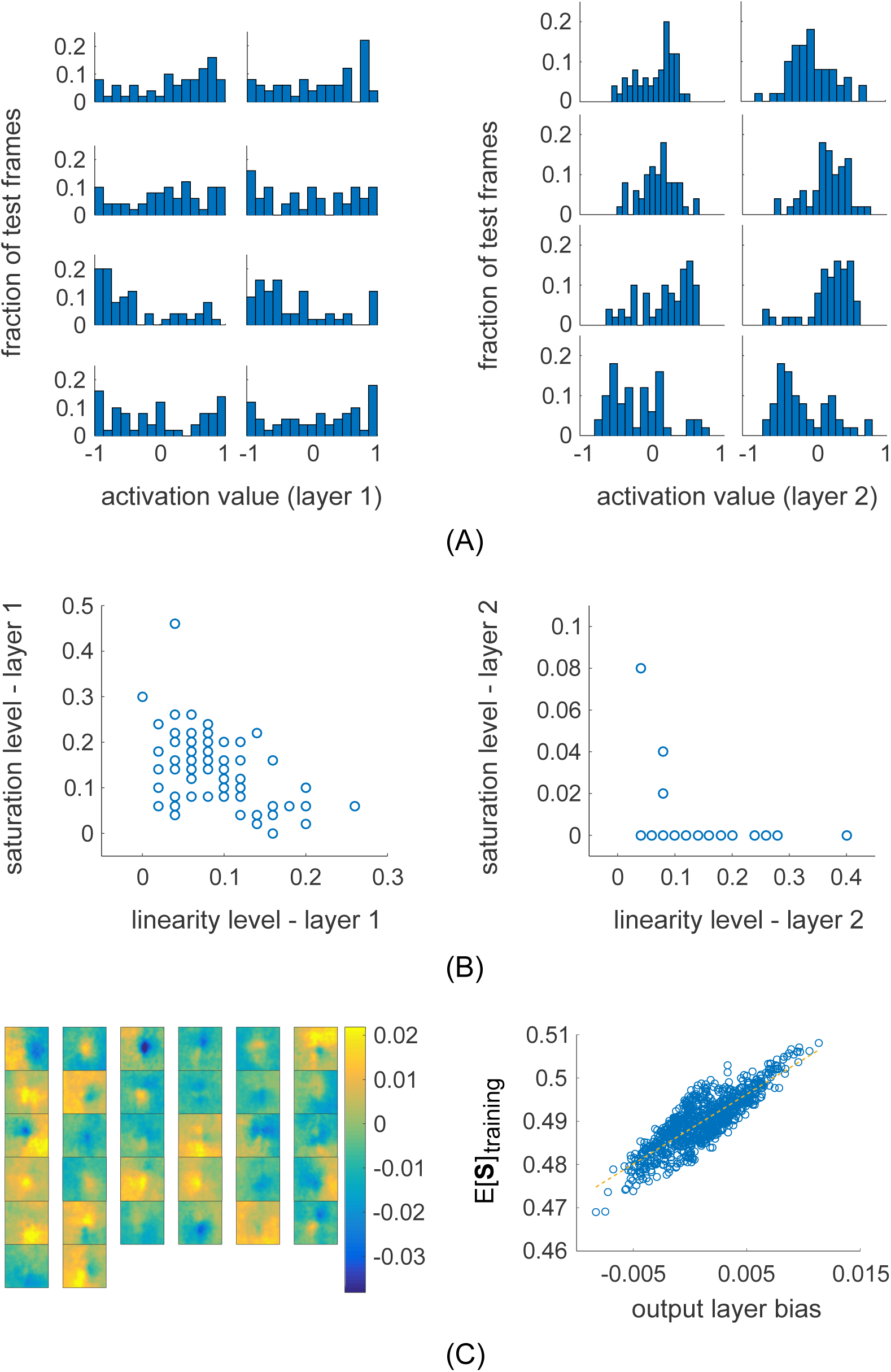
Non-linear decoding: analysis. **(A)** Activations of 8 example hidden units from the first (left) and second (right) hidden layer. **(B)** Saturation vs linearity index for the activations of units in the first (left) and second (right) hidden layers. **(C)** Weights (left) and biases (right) for the units in the output layer.

Furthermore, we performed some checks to verify that the network was successfully trained and that its performance was not hindered by operating in a suboptimal regime. First, we plotted the bias term of the output units against the average value of each corresponding pixel computed on the training dataset. It can be seen in Fig 5C that, as expected, these two quantities are directly proportional. Then, we computed the activation levels of all the units in the two hidden layers for each frame in the test dataset, to investigate the intermediate steps of the input-output mapping learnt by the network. Fig 5B shows examples of activation histograms for 8 units in the first (on the left) and second (on the right) hidden layer, selected at random. It can be observed that, especially for the first hidden layer units, the activation profiles are spread along the entire output range, that is between −1 and 1. Instead, the activation histograms of the second hidden layer units, while still covering the full range, are comparatively more concentrated around 0.

Such a result would seem to suggest that the transformation operated by the first hidden layer is more non-linear than that of the second layer, and it might help explaining why using a deeper network does not increase performance (S2 Fig). Indeed, when using sigmoid-like functions as non-linearities, there is the need to achieve a balance between the saturation – that is, when the output of a unit is heavily skewed towards the two extremes values – and the linear regime – that is, when the output of a unit mostly cluster around the linear part (an interval around 0) of the sigmoid-like function. This is because the former would hinder training by halting backpropagation at the unit, while the latter would fail to exploit the full representational power of the network. To quantify the interplay between these two different network behaviours, we computed a saturation and a linearity index for each hidden unity. They are defined as the percentage of test frames for which the absolute activation value is greater than 0.9 and less than 0.1, respectively. Results are shown in Fig 5B and appear to confirm that, while the second hidden layers behaves more linearly, the network overall operates in a balanced regime, away from extremely high saturation or linearity levels.

### Non-linear decoding does not outperform the optimal linear estimator

Quantitatively, the reconstruction accuracy of the ANN seems to have slightly outperformed that of the OLE. However, qualitatively, the predicted frames appear similar (Fig 6C). To check whether the performance improvements are significant, we performed a frame-wise comparison between the ANN and the OLE, which can be visualised in Fig 6A. As can be seen from the plot, most of the test frames cluster around the *y* = *x* line, with only a small subset of data points deviating from this trend in favour of increased ANN performance. Statistically, we quantified whether the two distributions were different by using a Wilcoxon signed rank test for the equality of medians. The resulting *p*-values were 0.17 and 0.62 for the MSE and the correlation coefficients, respectively, thus failing to disprove the null-hypothesis of equal performance between the ANN and the OLE.

**Fig 6.**
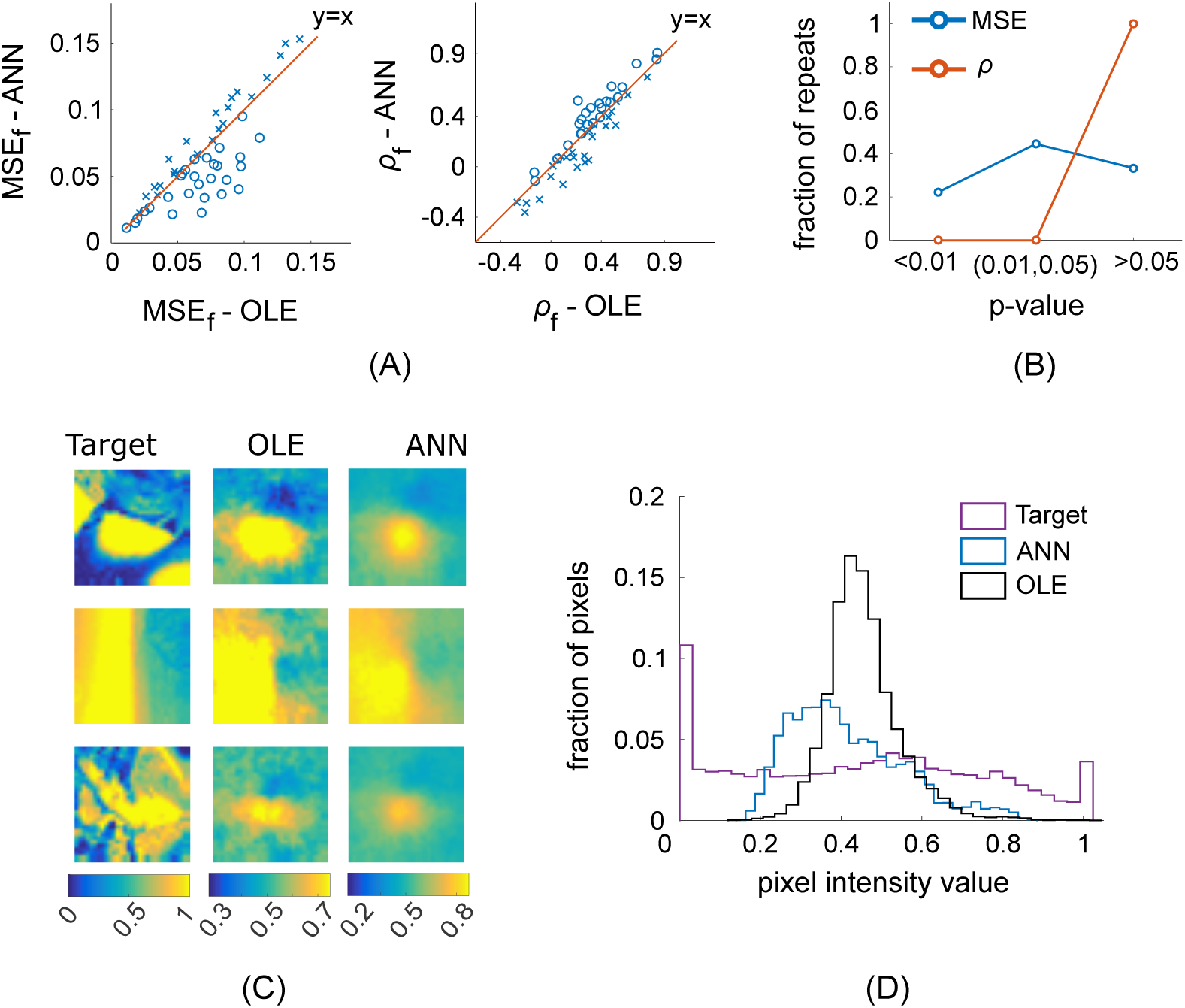
Linear vs non-linear decoding. **(A)** Performance comparison across frames (each data point is an individual image) for the MSE (on the left) and the correlation coefficient (on the right). The solid red line is the *y* = *x* line. Circles and crosses represent frames for which the ANN outperforms and underperforms the OLE, respectively. **(B)** Distribution of p-values across different network initialisations. The blue and red lines are for the MSE and the correlation coefficient, respectively. **(C)** Example of reconstructed frames using linear (middle column) and non-linear (right column) decoding. **(D)** Comparison of histograms of pixel intensity values. The target distribution is in purple, those generate with the OLE and the ANN are in black and blue, respectively.

To assess whether a different initialisation of the network parameters (which, in turn, could potentially result in the network converging on another local minimum) would have led to a different outcome, we repeated the same comparison analysis using 54 different copies of the same neural network architecture, each trained until convergence. The distributions of *p*-values from the Wilcoxon signed rank test across the network ensemble are shown in Fig 6B. While the differences in the correlation coefficient distributions are never statistically significant (not even at the *α* = 0.05 level), 20% of the ANNs significantly improved the MSE estimation (significance measured at the *α* = 0.01 level). It is possible that such an outcome derives from the ANNs being more effective at estimating the mean value of each frame, since this would have repercussion only on the MSE value and not on the correlation coefficient. An alternative hypothesis is that the lower MSE scores obtained by the ANN is a product of the differences in the distributions of pixel intensities for the reconstructions obtained with the ANN and the OLE. As shown in Fig 6D, the ANN-decoded distribution seems to have a less pronounced peak around the average value and to match the target distribution more closely. It is possible that, contrarily to the OLE, the presence of non-linearities allows the network to generate reconstructions with a pixel intensity distribution that more strongly deviates from a Gaussian.

## Discussion

This paper focused on pixel-wise stimulus reconstruction from 2-photon imaging recordings of mouse V1. First, we reconstructed the visual stimulus using the OLE and showed that a linear decoder already performs significantly better than chance, despite the overall performance level being relatively low. The pixel-wise algorithm performance was shown to be proportional to the level of coverage of the population receptive fields at each pixel. Then, the importance for decoding of pairwise neural correlations was assessed by comparing performance with that of a correlation-blind linear estimator (dOLE). We found that the differences in accuracy between the OLE and the dOLE are statistically significant on synthetic data, but not on recorded data. Finally, we compared the results obtained using linear and non-linear decoding and showed that, despite offering some improvements, especially in the estimated distribution of pixel intensities of the reconstructed frames, the non-linear algorithm did not significantly outperformed the OLE.

The near-equivalence between non-linear and linear reconstruction algorithms was not an obvious result. On the one hand, this might be related to the negligibility of pairwise correlations from a stimulus reconstruction perspective. On the other hand, Botella-Soler et al. report that their non-linear decoder, however used on retinal ganglion cells responses and complex but artificial movies, was only marginally affected by the removal of cell-cell noise correlations [14], but instead depended crucially on the spike history dependencies in the neural activity. In this work, instead, only the spatial pattern of responses was available for each stimulus frame, with no temporal information. It might be, then, that the ANN would have significantly outperformed the OLE had the full spatio-temporal pattern of responses been available. So far, there have been discordant results in the literature, with some studies showing an increase in performance with non-linear techniques [14–17], while others finding results more similar to this work [2, 18–20]. It is worth remarking, however, that, without computing the exact amount of stimulus information contained in the neural responses, any conclusion drawn from direct comparisons of linear and non-linear techniques is only valid with respect to the specific reconstruction algorithms and the neural responses recording methods that were employed. Indeed, it is possible that specific changes in the non-linear decoders could help increase the gap with linear decoders. For example, we could try incorporating a natural image prior to provide the reconstruction algorithms with knowledge about natural image statistics. Parthasarathy et al. [17] achieve this by enhancing the output of a linear decoder using an autoencoder-style neural network architecture, albeit applied to simulated responses from retinal ganglion cells. Another approach would be to experiment with different loss functions, since using the MSE usually produces highly blurred reconstructions by averaging over the full space of equally likely solutions [38]. Adding an additional term in the cost function that is able to capture a perceptual loss, like a distance in a pre-learned visual feature space [38], could be especially beneficial. However, such a perceptual loss should be preferentially tailored to the early visual system, rather than to high-level visual discrimination tasks [17].

The possibility of linear decoding being a general property of sensory neurons has been repeatedly conjectured [2, 3], strengthened by the observation that linear decoding is compatible with highly non-linear encoding strategies, and that it would allow downstream neurons to have access to various stimulus features by simple linear filtering of their inputs in the dendritic tree [2, 18]. Furthermore, through a combination of computational models and decoding, Warland et al. [18] observed that the ability of an ANN to outperform a linear estimator was not consistent, but instead depended on which specific non-linear neural code was used. Similarly, linear decoding was shown to be sufficient to extract most of the information contained in tonically firing neurons, but not in bursting ones [39]. Furthermore, evidence that linear algorithms are an adequate readout of neural activity supports the use of a fixed linear decoder when deriving efficient coding models of neural responses [40]. Finally, linear decoders have a simple and plausible biological implementation, although so would ANNs. Nonetheless, the generality of linear readout as a model of neural population decoding still remains open for discussion [3], with important theoretical and practical implications, for example in brain-machine interface devices. Related questions would be to investigate which features of the neural code are compatible with, or even favour, linear decoding [41], as well as under which experimental conditions non-linear algorithms might outperform linear ones, and whether the outcome can be predicted. In particular, it might be worth investigating whether the relative performance of linear and non-linear decoders on visual stimulus reconstruction depends on the specific combination of brain area (or cortical layer) under consideration and stimuli used [42]. For a full investigation of this topic, it will be essential to examine many different datasets, ideally through open source sharing of data and techniques.

Another question that is debated in the decoding literature is concerned with the role of pairwise neural correlations (specifically, noise correlations) on the reconstruction performance [6, 28, 33]. In this work, we found that neurons appeared to act as independent encoders when considering the experimental data, but not when analysing the synthetic data. This discrepancy does not seem to arise purely because of weak correlation levels, since it was consistent at all the levels of SNR explored. Instead, it might be due to the dissimilar dataset sizes, since results on synthetic data are not always robust to comparisons of subsets of 50 frames. Furthermore, given that the synthetic data was, by construction, devoid of noise correlations, which were instead present in the experimental data, we reasoned that such a difference might partly explain the observed results. Indeed, it has been shown that the interaction between signal and noise correlations has implications for decoding [28]. At the same time, it is worth remarking that being constrained to remove both signal and noise correlations at the same time is an important limitation of the current study. Hence, a more thorough investigation, such as repeating the same analysis using synthetic data generated with correlated noise (estimated from the experimental data), is needed to explore whether any difference between synthetic and recorded neural responses is due to the interplay of signal and noise correlations. Finally, another possible explanation is given by the pyramid wavelet model not being able to accurately represent pairwise neural correlations.

Full stimulus reconstruction from mouse V1 is, in general, a challenging task for many reasons, such as the low acuity of vision, and the high dimensionality of the stimulus to estimate. For example, the low acuity of mouse vision might explain why both decoders learned to reconstruct low resolution features better than high resolution ones. Additionally, some of the characteristics of this specific dataset, like a small number of training samples, the lack of fine-grained temporal information (more likely to happen with 2-photon imaging than with multi-electrode recordings), and a restricted number of imaged neurons, constituted other performance-limiting factors and might help explain the reasons behind the relatively low accuracy of both algorithms. Hence, improvements in 2-photon imaging conditions will potentially lead to better decoding performance. For example, recording a higher number of neurons would increase the receptive fields density at various pixels and results showed the direct proportionality between this density and the pixel-wise reconstruction accuracy. Furthermore, a higher number of neurons could also decrease the amount of training samples needed to reach certain performance levels [43].

At the same time, some studies also highlighted the importance heterogeneous tuning profiles when assessing the advantages of using larger neural populations for decoding [10, 44]. In the current population, however, the extent by which the receptive fields overlap indicates low heterogeneity, at least with respect to the receptive field location. Moreover, it is possible that recording from a larger population would also not be able to counteract the coarse-grained temporal properties of the responses [41], recorded at a low sampling rate (7.6 Hz) and averaged over approximately 500 ms. Indeed, it is likely that the lack of temporal information in the neural activity was a limitation not only to assess the relative accuracy of linear and non-linear algorithms, but also in terms of absolute performance levels. Furthermore, quantifying the amount of information lost with more aggressive time-averaging strategies could give insights into the different properties of temporal and rate codes in mouse V1 [7], from the perspective of a population of downstream neurons. Depending on how critical the role of temporal precision of the responses is, optical recordings, given their lower sampling rate, might be at a disadvantage with respect to multi-electrode arrays. Finally, an important avenue for future research is to extend the reconstruction to natural movies, to allow for a more complete investigation of decoding strategies in mouse V1.

## Materials and methods

### Data collection

The data were recorded by Antolik et al. [32] and released under the terms of the Creative Commons Attribution Licence. Briefly, the stimulus set is a collection of static images from David Attenborough’s BBC documentary *Life of Mammals.* Each image had 256 equally spaced luminance steps and was presented for 500ms, interleaved by 1474ms of blank grey screen. Recordings were made at 7.6Hz and putative spike counts were inferred using a deconvolution algorithm [45] for calcium traces. The average number of spikes across 5 consecutive 2-photon imaging frames was taken as the neuron response to a single stimulus frame. The final natural images used in the analysis are a patch of the displayed frames, centered around the location of the estimated receptive fields for the whole population, and down-sampled to 31 × 31 pixels. Details can be found in Antolik et al. [32]. Here, we used one of their image regions, containing 103 individual neurons. The training dataset, single-trial recordings, consists of 1800 image patches, while the test dataset, multi-trial recordings, has 50 image patches.

### Data analyses

#### Optimal Linear Estimator

We use a multi-input, multi-output linear estimator to decode the stimulus from the neural responses [19, 23], that is

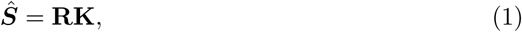

where *Ŝ* ∈ ℝ^*T*×*N_p_*^ is the reconstructed stimulus, **K** ∈ ℝ^(*N_n_*+1)×*N_p_*^ is the matrix of the linear filters and **R** ∈ ℝ^*T*×(*N_n_*+1)^ is the neural responses matrix. Here, *T* is the number of training frames, *N_p_* is the number of pixels in each frame and *N_n_* is the total number of neurons. The extra column in the **R** matrix correspond to the bias term, i.e. it is a column with entries all equal to 1. Finally, the data in **R** is normalised so that each column, excluding the last one, has 0 mean and unit variance. We estimate the optimal linear decoding filters by minimising the reconstruction MSE with L_2_ regularisation, with weight given by λ. The solution is given by the Optimal Linear Estimator (OLE) [19]:

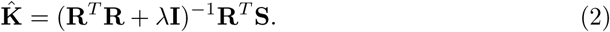

The entire dataset was divided into training and test data; **K̂** was computed using only the training dataset, while performance is measured on the test dataset. A 5-fold cross validation was used to find the optimal value of λ.

To quantify the dependency of the performance on the number of neurons, sampling without replacement was used to select subsets of different sizes, from 1 to 102, from the pool of all available neurons. For each size, we selected 103 different subsets (to match the number of subsets of size 1), applied the OLE to each of them and computed the mean and the standard deviation of the performance across repetitions. The results was an empirical curve establishing the relationship between number of cells and reconstruction performance. Subsequently, such a curve was fitted with a function given by the ratio of two quadratic polynomials, that is:

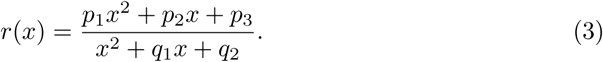

The value of the coefficient *p*_1_ would then give the expected performance in the limit of the number of neurons going to infinity. Using the ratio of linear polynomials gave similar results.

#### Diagonal Optimal Linear Estimator

To investigate the role of correlations, we built a correlation-blind decoder and quantified its impact on the reconstruction performance [6, 28, 33]. Specifically, the off-diagonal elements of the matrix **R^*T*^R** were set to 0, since, by construction, given that the neural responses have been standardised, they contain all the information about pairwise correlations between neurons. Thus, the diagonal OLE, or dOLE, is defined by:

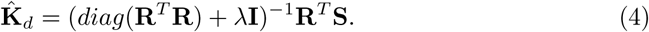

As with the OLE, we optimised the linear filters on the training dataset using Equation 4 and evaluated performance on the test dataset.

#### Fully connected neural network

To perform non-linear stimulus reconstruction, we used a 2-layer fully connected neural network with 64 and 32 hidden units, respectively. Hidden units have the hyperbolic tangent as their activation function, while a sigmoid with gain of 10 was used for the units in the output layer. We used dropout [46] (with probabilities 0.5 and 0.4 for the first and second hidden layer, respectively) and batch normalization [47] to regularize the network and prevent overfitting. The described hyper-parameters were obtained using a 18-fold cross-validation on the training dataset. The network was trained end-to-end for 201 epochs using the backpropagation algorithm and Adadelta update scheme. For the first 101 epochs, the default Adadelta learning rate of 1 was used, and it was decreased to 0.001 afterwards. Binary cross entropy was used as the cost function to train the network. We trained 54 different instances of the same architecture and selected the final one using performance computed on a validation dataset (approximately 10% of the training dataset). Finally, saturation and linearity indices were computed for the activation profiles of each hidden unity. They are defined as the percentage of test frames for which the absolute activation value is greater than 0.9 and less than 0.1, respectively.

#### Quantification of performance

Performance was quantified using two commonly employed measures [48], i.e. mean squared error (MSE) and Pearson correlation coefficient between each target frame and its reconstruction. Since a frame **s** is defined as a 1-dimensional vector of pixels {*s_p_*}_*p*∈𝓟_ with length equal to the number of pixels *N_P_*, the MSE is given by:

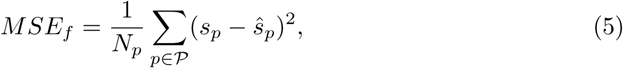

where *s_p_* is a single pixel in a target frame, and *ŝ_p_* is the corresponding pixel in the reconstructed frame. The median value over all frames in the test dataset gives the performance measure 〈*MSE_f_*〉. The Pearson correlation coefficient is defined as:

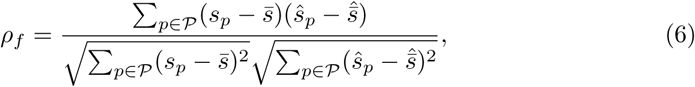

where *s̄* is the average pixel value of the target frame and *s̄̂* is the average pixel value of the reconstructed frames. The median value of the above quantity over the test dataset gives the performance measure denoted by 〈*ρ_f_*〉.

To quantify how well we can estimate individual pixels, i.e. specific spatial locations, we used two analogous measures, *MSE_p_* and *ρ_p_*. These are defined similarly to Eq (5) and (6), but with sums and averages computed over all frames in the test dataset, separately for each pixel.

#### Statistical tests

To assess the statistical significance of the reconstructions with various decoders, we characterized the chance level as the performance of the same algorithm applied to data in which the neural responses had been randomly shuffled to remove the input-output relationship between visual stimuli and neural activity. Specifically, each spatial pattern of neural responses for the *i*–th frame was randomly assigned to a different frame, *j*. This allowed us to construct a null hypothesis scenario of decoding performance purely due to chance. Such a shuffling was repeated 200 times, and chance-level performance values (the shuffled distribution) obtained as the average over all repetitions. We then assessed statistical significance using a paired Wilcoxon signed rank test between the original and the shuffled distribution of accuracy (AB test 1). When needed, we used the Holm-Bonferroni correction to control for multiple comparisons. For completeness, we also performed a second statistical test (AB test 2). Indeed, we can consider each shuffling repetition as an individual trial Bernoulli process where success is defined as obtaining a median performance value over the test dataset worse than that computed on non-shuffled neural responses. If results are only due to chance, success probability is *p* =0.5. Hence, we can count the number of positive outcomes over the 200 repetitions and then calculate the *p*-value as the probability of observing at least as many successes under the null hypothesis of a chance-level decoder. When needed, the Holm-Bonferroni correction was used.

#### Pyramid Wavelet Model

A pyramid wavelet model was used to characterize the neural responses. Such a model relies on a non-linear transform *L*(·) of the stimulus to reach a feature space that has a linear relationship with the neural activity. Specifically, the pyramid wavelet model used in this study is defined by a set of base wavelets, and a non-linear transform from pixel space to wavelet space. For the former, we used a set of cosine (even) and sine (odd) Gabor filters with different frequencies, locations, phases and orientations, while the transform is given by a linear projection of a stimulus frame into each wavelet followed by a point non-linearity (either a ramp function or the sum of squares from quadrature-phase wavelets, to model both simple and complex cells). The result is a vector of wavelet coefficients *L*(**s**), whose dimension if given by the number of features (denoted by *N_F_*). Then, the response *r_ij_* of an individual neuron *j* to a single stimulus frame *i* is given by *r_ij_* = *L*(**s**^*i*^)**h**^*j*^. Here, **h**^*j*^ is a weight vector, specific to each neuron and with size given by the dimension of the feature space, whose entry 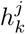 quantify how much the response of neuron *j* depends on feature *k*. Such a vector was estimated using regularized linear regression, specifically the *L*_2_*boost* algorithm, to encourage sparseness [32, 37]. We used the open source STRFLab toolbox [49] to fit the model.

We used 6 different orientations (*θ* = *kπ*/6, *k* = 0, 1, 2, 3, 4, 5), and 5 different spatial frequencies on a logarithmic scale; for each frequency *f*, the standard deviation of the Gaussian envelope was given by 2*πσ* = *κ*1/*f*, where 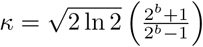 [50], and *b* is the frequency bandwidth in octaves (here, *b* = 1.5). Furthermore, 5 phases were considered, that is the negative and the positive part of each half-rectified even and odd wavelet, together with the phase-invariant projection given by the sum of squares of two quadrature phase wavelets [25, 26, 32, 37]. Finally, the aspect ratio was fixed to 1, and, at each orientation and scale, adjacent wavelets were separated by a fixed number of pixels corresponding to the standard deviation of their Gaussian envelope. The total number of wavelets used is 33990.

#### Receptive field coverage

To obtain the number of receptive fields at a given pixel, we derived a linearized version of each neuron’s receptive field. First, we summed all the base wavelets used in the pyramid Wavelet Model, each multiplied by its respective entry in the weights vector **h** (detailed in the pyramid Wavelet model section). Such a procedure computed a different weighted sum for each neuron. A single elliptical Gabor function was then fit to the result, and the Gaussian envelope taken as the boundary of each neuron’s receptive field. If the fit failed, the Gaussian envelope of the base wavelet corresponding to the highest entry in the weights vector **h** of the pyramid Wavelet model was used.

#### Surrogate data generation

Synthetic neural response were generated using the parameters estimated by the pyramid wavelet model. First, we produced a new and larger natural image dataset (the synthetic dataset) using data from both the VanHateren dataset [51] and the Berkeley segmentation dataset [52]. After resizing all frames to 256 × 256, we extracted the 128 × 128 central patch, down-sampled it to a resolution of 31 × 31, and finally used contrast normalization. The pixel intensities distribution of the synthetic dataset was matched to that of the original stimulus dataset. We augmented the data by applying three transformation to each frame (vertical and horizontal flipping, and reverse contrast). The final length of the dataset was of 18668 frames. Examples of stimuli and neural responses can be seen in S1 Fig. Finally, we added uncorrelated Gaussian noise at various level of SNR (between 0 and 200) and set to 0 any negative firing rates.

## Supporting information

**S1 Fig.**
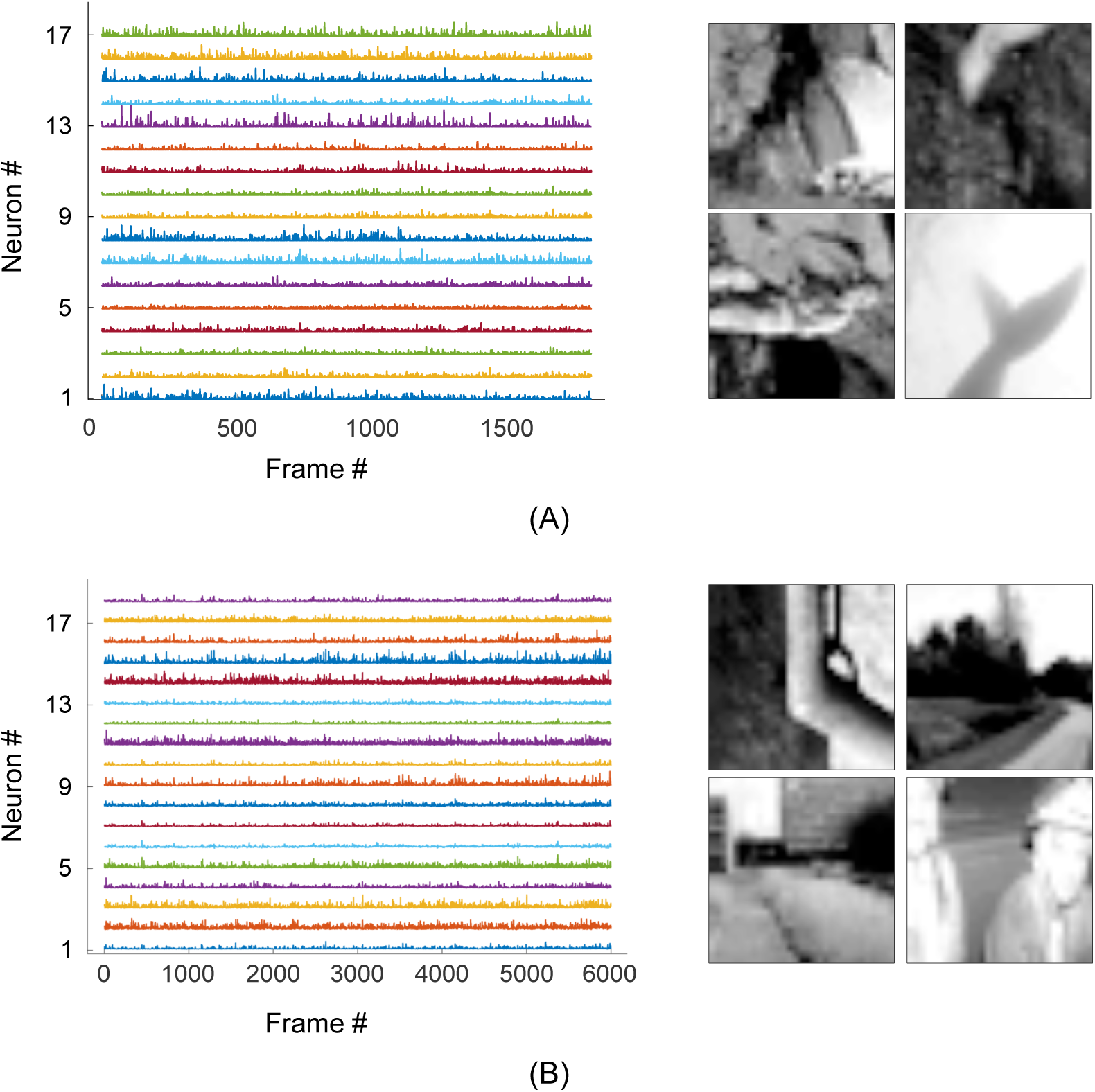
Samples from the datasets. (A) Example neural responses (left) and natural stimuli (right) from the experimental dataset. (B) Example neural responses (right) and natural stimuli from the synthetic dataset. The neuron id numbers do not refer to the same cells as in (A).

**S2 Fig.**
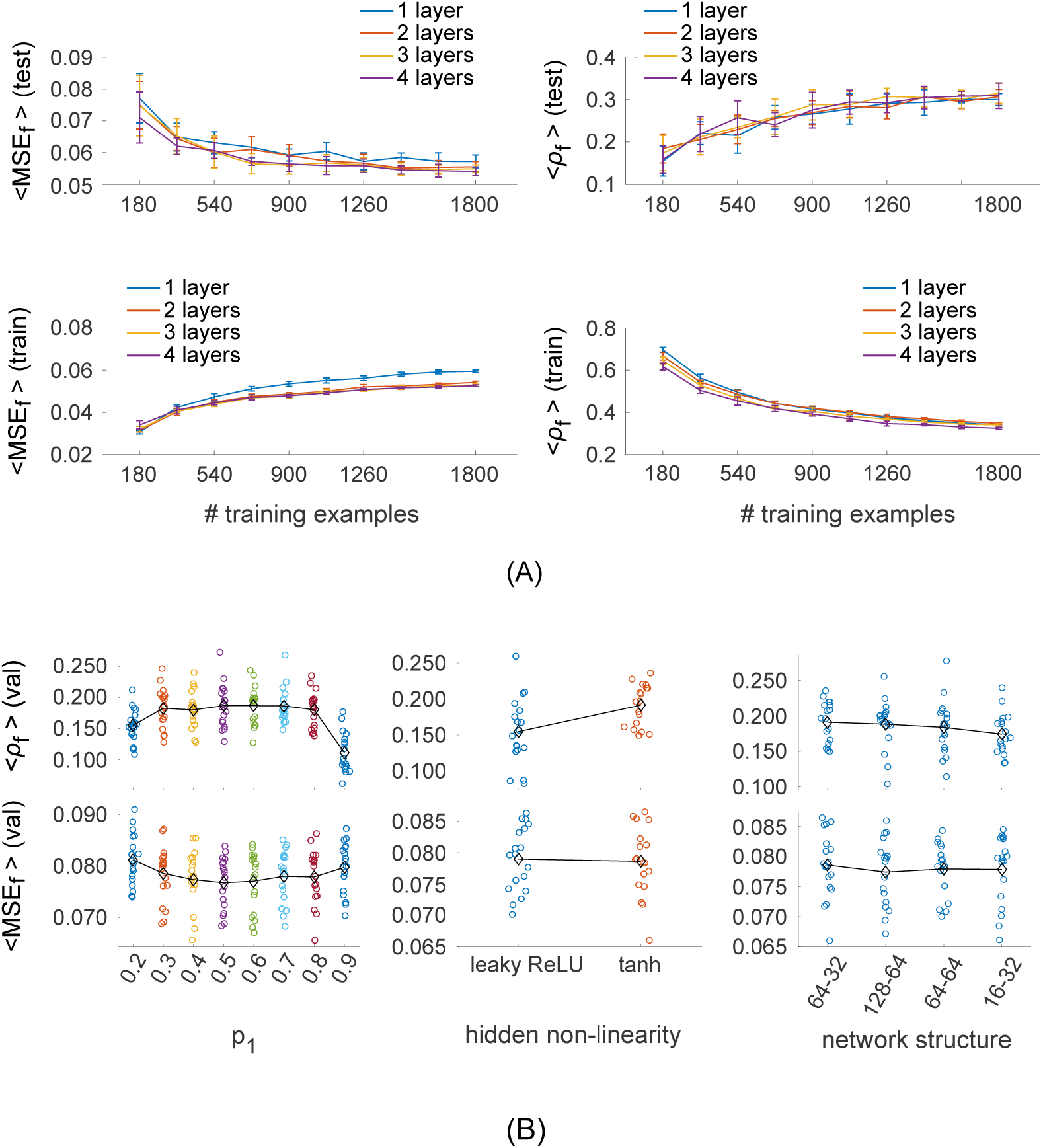
Exploration of the ANN hyper-parameters space. (A) ANN performance vs the number of layers and the number of training samples used. We trained several networks with number of hidden layers varied between 1 and 4 (the number of hidden units in each layer *n_l_* was equal to 2^*n_l_*+4^). The number of training samples was varied between 180 and 1800, with steps of 180 frames. For each combination we trained 10 different networks, each using a different random subset (of fixed size) of the training dataset. Results are reported using the average and the standard deviation over those 10 repetitions. (B) ANN performance vs. the dropout probability (left), the hidden non-linearity (middle) and the number of units in the two hidden layers (right). In the left-most plot, the variable *p*_1_ is the dropout probability for the first hidden layer. The dropout probability for the second hidden layer, *p*_2_ was set at *p*_2_ = *p*_1_ – 0.1. In the right-most plot, each network structure is described by a pair of numbers, indicating the number of hidden units in the first and second layer, in this order.

**S3 Fig.**
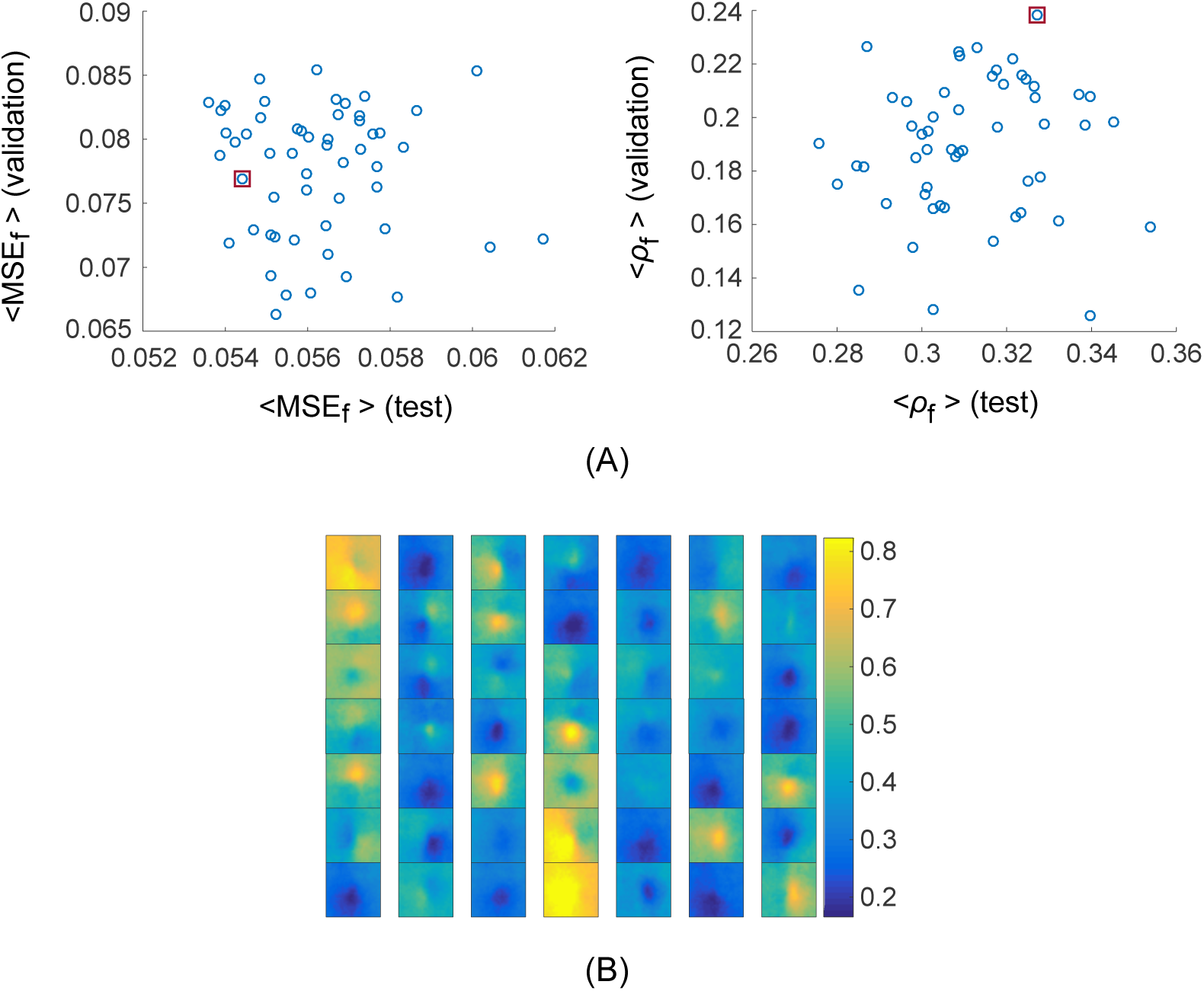
ANN ensemble performance and reconstructed frames. (A) Test versus validation performance (MSE on the left and correlation coefficient on the right) for each of the initialisation of the final ANN architecture. The data point enclosed by a red square is the model selected for further analysis. (B) Reconstructions produced by the ANN for 49 out of 50 frames in the test dataset. All images share the same colourmap.

**S1 Appendix. Multi-target regressor stacking.** Multi-target regressor stacking (MTRS) [53] is a multi-steps process. The first is equivalent to the OLE, that is we fit a series of single-output linear regressions models with L_2_ regularisation. Then, the original input matrix and the target variable predictions from the previous step are concatenated to form an augmented input matrix. Finally, a new, augmented, linear regression model is estimated using the augmented input matrix. Novel predictions are made following the same procedure. The underlying principle is that including information about the predicted output allows the model in the second stage to consider correlations between the target variables. The median performance values obtained using this reconstruction algorithm were 0.0684 and 0.282 for the MSE and the correlation coefficient, respectively.

## Acknowledgments

We thank Wilten Nicola and Antonia Creswell for many useful discussions.

## Authors Contribution

Conceived and designed the research and the analysis: SG AAB SRS. Performed research: SG. Processed data: SG. Analyzed the data: SG. Contributed reagents/materials/analysis tools: SG AAB SRS. Wrote the paper: SG AAB SRS. Discussed the data and commented on the manuscript: SG AAB SRS. AAB and SRS are Joint Senior Authors.

